# Community Web Portal for Open Collaboration in the Martini Force Field Initiative

**DOI:** 10.64898/2026.01.27.701988

**Authors:** Daniel P. Ramirez-Echemendia, Luís Borges-Araújo, Chelsea M. Brown, Riccardo Alessandri, Siewert J. Marrink, Paulo C. T. Souza, D. Peter Tieleman

## Abstract

The Martini coarse-grained force field is widely used for biomolecular simulations by a large and rapidly expanding community worldwide. Over time, the development of Martini parameters, tools, and documentation has become increasingly dispersed across numerous research groups, leading to fragmentation and making it challenging for users and developers to keep track of the latest models, software, and best practices. Consequently, the development of Martini as a genuinely community-driven process has grown into a bottleneck. In response, the Martini Force Field Initiative (MFFI) has been established as an open-science effort to coordinate and support the collaborative development of all Martini resources. Here, we introduce the MFFI web portal, a platform designed around five core pillars: (i) avoiding reliance on a single group or local server; (ii) minimizing long-term maintenance overhead; (iii) reducing technical barriers for contributions; (iv) providing a unified home for parameters, tools, tutorials, example workflows, and research outputs; and (v) enabling timely dissemination of updates to the community. To achieve this, we use Quarto to generate a static website authored in Markdown, lowering the technical barrier to making contributions, and serverless architectures on Amazon Web Services for scalable, event-triggered backend operations. The source code is hosted in a public GitHub repository under an MIT license, with automated deployment via GitHub Actions and a contribution model based on pull requests for quality control. This design creates a sustainable, low-maintenance, and collaborative infrastructure that consolidates Martini resources and supports transparency. More broadly, our design exemplifies a transferable pattern for building open, community-oriented platforms for molecular modeling and computational science.

## 1 Introduction

Over the past two decades, coarse-grained (CG) molecular dynamics has emerged as a central tool for examining molecular systems at spatial and temporal scales often beyond the reach of atomistic simulations [1, 2]. In CG models, groups of atoms are represented as effective interaction sites, trading atomic detail for substantial gains in computational efficiency while preserving the essential physico-chemical properties of the systems. Within this landscape, the Martini CG force field (FF) [3, 4] has become one of the most widely adopted CG frameworks for (bio)molecular simulations [5], spanning applications from lipid membranes [6] and proteins [7] to nucleic acids [8], carbohydrates [9], polymers [10] and nanomaterials [11, 12]. Its general-purpose nature, combined with its large library of existing parameters and tools, has led to its adoption by a broad, interdisciplinary community of users and developers worldwide.

As a result of this rapidly expanding community, Martini development has naturally transitioned from a closely coordinated effort among a few core research groups to a more distributed network involving numerous contributors [4, 13]. New parameter sets, extensions to novel chemical classes, specialized tools for system building and analysis, and domain-specific workflows are now routinely developed by groups with diverse scientific goals and computational practices. Although this shift towards broader participation was intentionally pursued and has brought clear scientific benefits to the project, it has also revealed significant challenges in coordination and sustainability. Martini-related resources are currently scattered across personal and institutional web pages, GitHub repositories, supplementary materials of individual publications, and various community channels. Thus, it can be difficult for users, especially newcomers, to identify which parameter sets are current and recommended, which tools are actively maintained, or where to find authoritative guidance on model selection or suggested validation. For developers, duplication of effort, inconsistent documentation, and lack of a clear “home” for new contributions can slow down the dissemination of advances and large-scale user testing. Moreover, many of these existing resources depend on a single research group or local servers, making them vulnerable to personnel turnover and evolving institutional priorities.

These issues are not unique to Martini. Similar problems have also been recognized in many areas of computational science where widely used methods and models emerge from academic research groups. In response, the broader open-science movement has emphasized principles and practices designed to ensure that scientific resources remain findable, accessible, interoperable, and reusable (FAIR) [14–16]. Notably, some community-oriented initiatives in molecular modeling have already worked on implementing these principles, namely PLUMED-NEST [17], NMRlipids [18], OpenMM [19, 20], and Open Force Field [21]. As such, a FF as widely used and continuously evolving as Martini stands to benefit strongly from a similar framework that supports open, collaborative development and transparent dissemination of models, tools, and knowledge. The Martini Force Field Initiative (MFFI) has been established to address this need by formalizing Martini as an open-science effort rather than a collection of loosely connected group activities. A central objective of MFFI is to provide an infrastructure through which the community can coordinate model development, share tools and workflows, and converge on best practices, while preserving the flexibility and creativity that have characterized Martini’s growth. Realizing this objective requires more than simply hosting files on a central server, it calls for a platform that reflects the realities of academic software development (e.g., high turnover in personnel, heterogeneous technical expertise, and limited time and resources for long-term maintenance) while aligning with contemporary practices in open, reproducible science.

As part of this effort, we started by outlining the set of practical requirements that the MFFI platform needed to satisfy. First, it should not rely on a single group, institution, or local server as its operational backbone, thereby avoiding single points of failure and easing the transition when core developers move or change roles. Second, long-term maintenance overhead must be minimized: routine operations such as deployment, scaling, and basic security should be automated as much as possible, recognizing that much of the day-to-day work will be carried out by PhD students and postdoctoral researchers whose involvement is time-limited. Third, it must present a low technical barrier to contribution, allowing domain experts in biomolecular modeling, who may not be web developers, to contribute content using simple workflows. Fourth, it should act as an entry point, or “home”, to the Martini community, organizing parameters, tools, tutorials, example workflows, and research outputs in a structure that makes it easy for users and developers to find relevant resources. Finally, it should support timely dissemination of updates, enabling the community to be notified of new models, tools, and documentation and thereby reducing the lag between development and adoption.

In this work, we introduce the MFFI web portal, our implementation of a platform designed around the five pillars mentioned before. At the operational level, we employ serverless backend architectures on Amazon Web Services (AWS) to provide scalable and robust infrastructure without the need for dedicated local servers or system administration [22, 23]. The portal’s front end is implemented as a static site generated using Quarto [24], with content authored in Markdown. This choice allows contributors to work with a human-readable, widely used text format that integrates naturally with version control, whereas Quarto’s literate-programming features facilitate the inclusion of executable examples, code snippets, and reproducible workflows. The entire site is maintained in a public GitHub repository under a MIT license, reflecting the open-source ethos of MFFI and making both content and infrastructure transparent and reusable [25]. Automated deployment via GitHub Actions ensures that updates merged into the main branch are reliably propagated to the live site with zero manual intervention [26], while a contribution model based on pull requests provides a mechanism for quality control, discussion, and review [27]. Importantly, this platform fully replaces an earlier version of the portal historically hosted at https://cgmartini.nl, which, despite the limitations discussed above, supported the community for many years in adopting and developing Martini. For this reason, we decided to preserve the previously established content organization and high-level layout to ensure a smooth transition.

Beyond addressing immediate organizational challenges for the Martini community, the MFFI web portal is intended as a concrete example of how modern web technologies, open-science practices, and cloud-based infrastructure can be combined to support community-driven development of scientific models and tools. Although the specifics of the content (e.g., FF parameter sets for CG molecular simulations and workflows for building and analyzing large systems) are particular to Martini, the underlying design principles are general. The approach we describe is readily transferable to other communities and computational frameworks facing similar issues.

## 2 Results

### 2.1 Web Portal as a unified entry point to Martini

The MFFI web portal (https://cgmartini.nl) serves as the main entry point to the Martini community and was designed to present all resources in an easily discoverable way. The landing page provides a concise overview of the Martini FF, highlights the most recent news, and points users to key review and methods papers. In addition, a persistent navigation bar exposes the different content categories: *Tutorials, Downloads, Publications, Announcements*, and *Contact*, alongside a *Help* menu linking to community support channels.

These categories correspond with the most commonly needed Martini resources for every-day use (Fig. **1**). *Tutorials* provide step-by-step workflows and teaching material for setting up, running, and analyzing Martini simulations; *Downloads* centralize FF parameter files and example inputs, organized by version and molecular class; *Publications* curate methodological and application papers that make use of the Martini FF; *Announcements* serve as a community news feed, directly linked to an email delivery system that ensures timely notifications when new content is released; *Contact* outlines a list of all Martini contributors with contact information and a form for general inquiries to the team; and *Help* features a Frequently Asked Questions (FAQ) section, and links to a GitHub issue tracker and discussion board. Implementation details are described in the Methods section.

**Figure 1.**
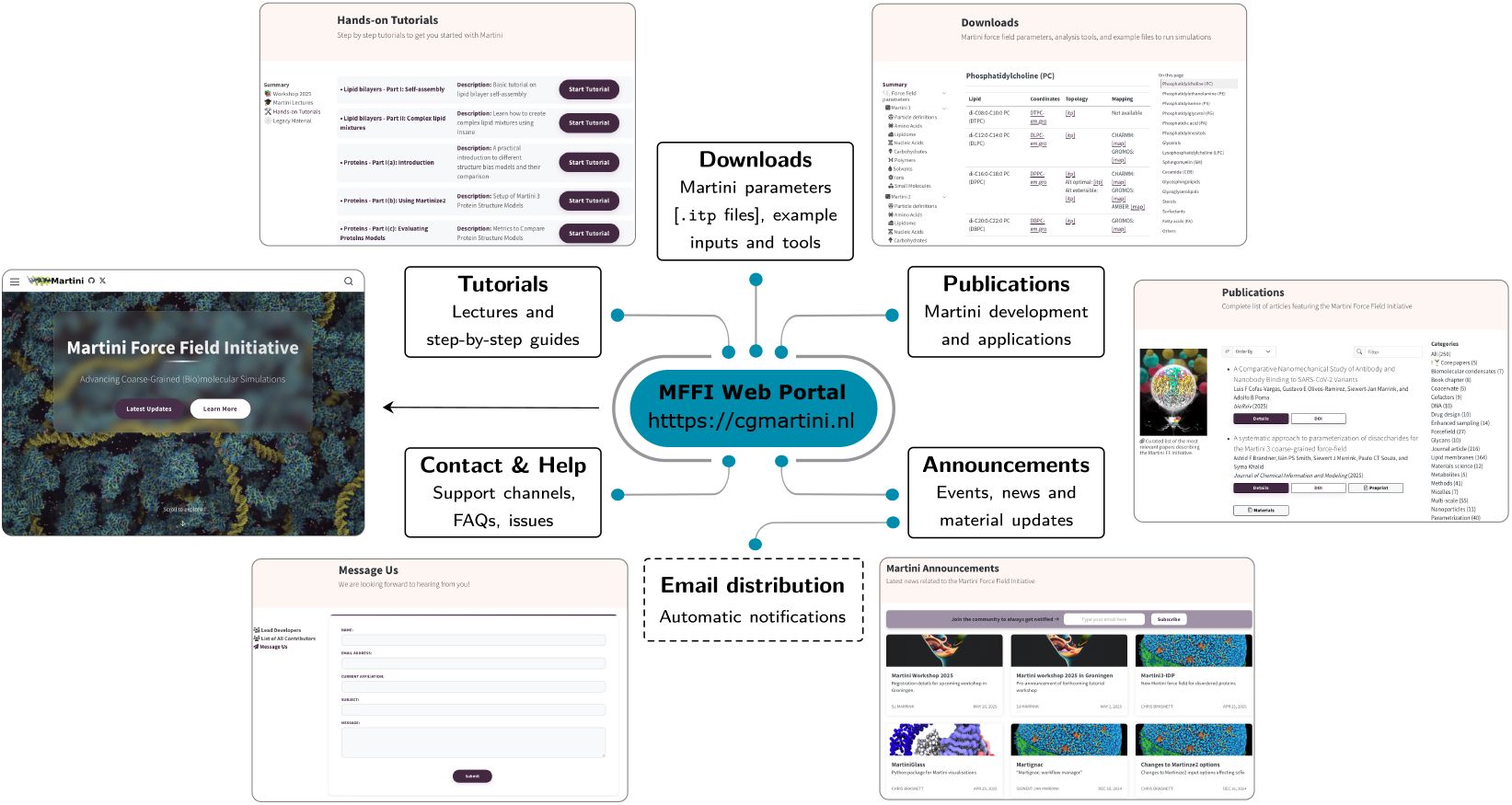
Schematic overview of content organization in the MFFI web portal. The portal is organized around the following resource types: *Tutorials, Downloads, Publications, Announcements*, and community support (via *Contact* and *Help*). New announcement posts are distributed to subscribed users via email notifications.

### 2.2 Lowering barriers to community contributions

A key design objective of the MFFI portal is to make community contributions routine rather than treating them as exceptional events, by using workflows that are already standard in open science development guidelines (Fig. **2**). Content is authored using Markdown syntax, version controlled in Git, and submitted via pull requests in GitHub. This “docs-as-code” model using a widely adopted markup language such as Markdown lowers the technical barrier to contribution for experts on different domains (e.g., FF developers, software engineers, and tutorial authors), by eliminating the need for prior expertise with custom web development platforms. At the same time, it preserves quality control through a lightweight review process: proposed changes are attributable to specific contributors, can be discussed inline, and are auditable through an explicit history of revisions. As a result, portal updates become citable artifacts, with clear authorship and a persistent record of decisions, similar to how code changes are managed in mature open-source projects. The repository contribution guide further streamlines this process by providing a list of supported contribution types (publications, announcements, tutorials, tools, FF parameter) and a local preview workflow for contributors to validate changes before submission (see Supplementary Information for details).

**Figure 2.**
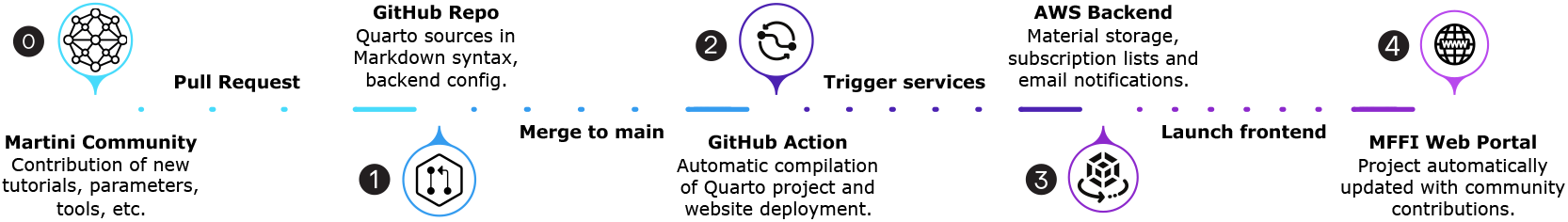
Contribution and deployment workflow. Community members submit contributions via pull requests to the public GitHub repository. Upon approval and merging, automated workflows regenerate and deploy the static front end and synchronize announcement content to backend services using GitHub Actions, ensuring reproducibility.

This contribution model is tightly coupled to the portal’s role as a centralized hub for parameters, tutorials, and tools. The portal organizes resources to minimize navigation depth and reduce ambiguity about where to find current, expert-endorsed Martini resources. The *Downloads* page groups files by Martini version and molecular class, helping users rapidly identify the appropriate FF parameter sets and reducing the inadvertent use of outdated files. In parallel, *Tutorials* and *Tools* provide practical workflows and software around common user tasks (e.g., topology/structure generation, multiscale setup, and analysis), translating expert knowledge into reusable, task-oriented points for both beginners and experienced users alike. Together, these features turn the portal into a living catalog of resources and a maintained record of how those resources evolve over time.

### 2.3 Decentralized maintenance and low operational overhead

Since its initial deployment, the MFFI web portal has operated without a dedicated, continuously administered server. Routine updates are handled through standard GitHub workflows (issues, discussions, and pull requests), and are automatically propagated to the live site: whenever a pull request is merged into the `main` branch, a GitHub Action renders the Quarto project and deploys the updated portal without manual intervention (Fig. **2**). This level of automation makes decentralized maintenance practical. As a result, dependence on any single individual or local administrative environment is reduced and responsibilities can transition cleanly as roles rotate.

On the backend side, the operational footprint is also minimized through serverless event-triggered architectures, in which services are invoked only in response to specific actions rather than maintained as a continuously running infrastructure. Consequently, day-to-day management is largely limited to minor editorial tasks (e.g., reviewing incoming contributions, enforcing consistent metadata and formatting, and maintaining cross-references and links). As such, the platform remains responsive to new resources while remaining robust to personnel turnover.

### 2.4 Community communication and dissemination of updates

The *Contact* section of the portal offers an up-to-date directory of contributors to the Martini FF, including affiliations and contact details. This makes the “who to ask” problem tractable for users, reducing reliance on informal knowledge, or third-party information discovery. Additionally, a general contact form routes general inquiries (e.g., collaboration proposals) to the MFFI team for triage and follow-up (Fig. **3-A**). The Help section links to a GitHub discussions board and an issue tracker, providing transparent channels for questions, feature requests, and bug reports. Together, these pathways promote reproducible support via searchable threads, enable moderation and prioritization, and create a durable record of resolutions that can benefit future users. By making these contact and support mechanisms visible and standardized, the portal facilitates scientific exchange and helps align user needs with ongoing development efforts.

**Figure 3.**
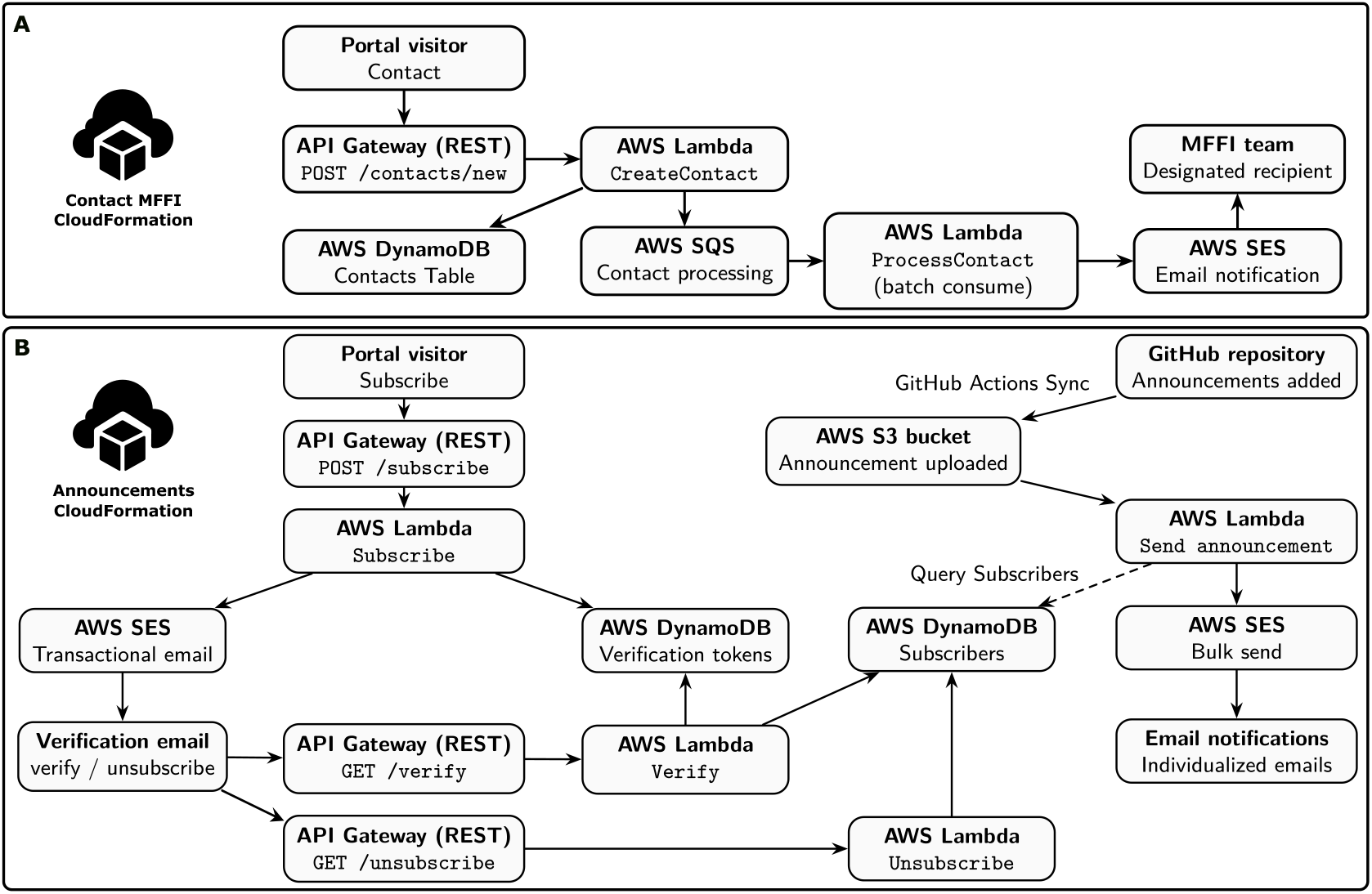
Serverless backend architectures supporting portal communication services. **(A)** *ContactsStack* handles contact-form submissions and asynchronous forwarding to the MFFI team for triage and follow-up. **(B)** *AnnouncementsStack* manages subscriptions, including email verification; and delivers announcement notifications.

Community members can easily create announcement posts to promote events, new parameter releases, tools, tutorials, workshops, and relevant publications. This broadens dissemination and reduces the delay between producing a resource and its adoption by others, shifting communication from a fragmented “pull” model to a proactive “push” model. Announcements are presented as a persistent portal feed and are coupled with an opt-in email notification system that alerts subscribers to new posts (Fig. **3-B**). As described in Methods, posts and email notifications are generated from a shared source of truth, ensuring consistent dissemination across channels with minimal duplication.

### 2.5 Martini Workshop 2025: rapid, low-friction collaboration

The 2025 Martini Workshop in Groningen, The Netherlands (August 10–15, 2025) served as a realistic stress test for collaborative content production on a fixed timeline (Fig. **4**). Over the course of this five-day event, instructors and developers from multiple groups worldwide contributed workshop material spanning lectures, hands-on tutorials, tool descriptions, and example inputs. The contribution workflow remained effective across authors with heterogeneous technical backgrounds, who were able to focus on scientific and pedagogical quality, while maintainers applied minor edits to enforce consistent structure and formatting. In addition, the workshop underscored the benefits of treating training material as version-controlled objects. As multiple contributors iteratively refined content, the workshop collection improved in clarity and completeness. The rapid pull-request–based corrections (e.g., commands, file paths, dependencies, and expected outputs) made also possible quick real-time fixes during the event. Thus, the materials resulting of this event transitioned from single-time resources to durable, reusable, and citable community assets.

**Figure 4.**
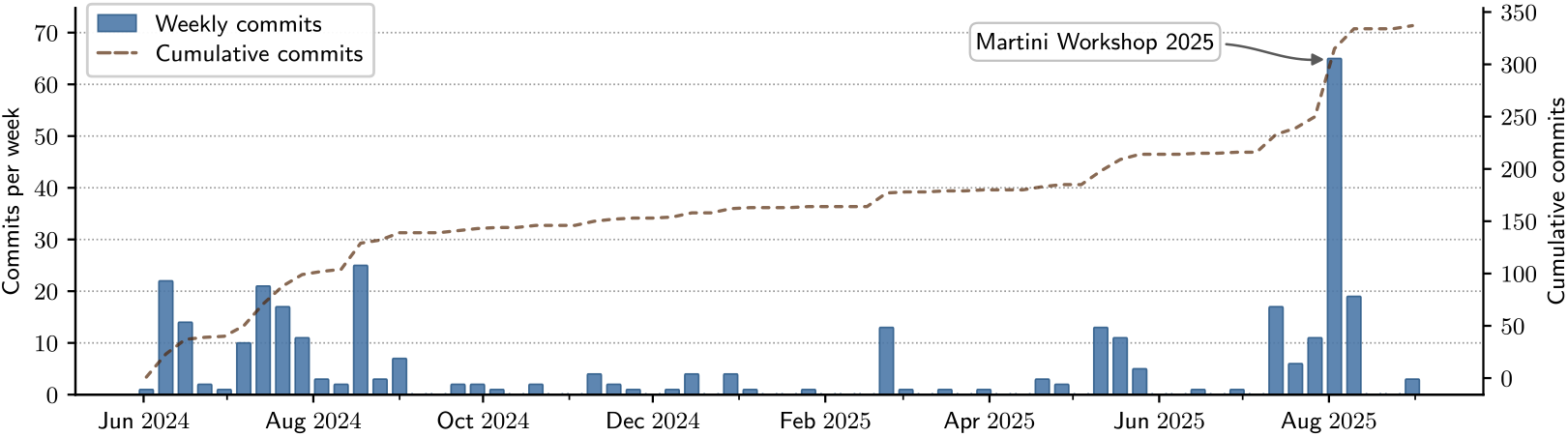
GitHub activity timeline for the MFFI web portal repository. Weekly commit counts are shown as bars and cumulative number as a dashed line (right axis), summarizing the growth of repository contributions over time. The highlighted peak in the first week of August 2025 coincides with the Martini Workshop 2025 in Groningen.

### 2.6 Generalization and transferability beyond Martini

Although this portal was developed for Martini, its architecture is intentionally modular and largely domain-agnostic. Martini-specific aspects are concentrated in the content layer, including domain terminology and the taxonomy of resources (e.g., parameter/topology formats and organization practices). In contrast, the operational mechanisms are generic: community contributions are mediated through GitHub pull requests and review, the front end is built and deployed automatically via GitHub Actions, and portal functionality is limited to a small set of serverless backend primitives (contact handling, subscription verification, and announcement-triggered email delivery). This separation of concerns provides a practical map for other computational communities that face similar constraints.

## 3 Discussion and Conclusions

The MFFI web portal was developed in response to a practical challenge faced by mature computational ecosystems: as community scales, infrastructure can become a limiting factor for coordination, discoverability, and sustained collaborative development. In Martini, growth in parameters, tools, and educational material has been accompanied by increasing fragmentation across group websites, supplementary files, and independent repositories, which creates friction for users attempting to identify current resources and slows down the feedback loop that enables community vetting and iteration. The MFFI portal targets this bottleneck by coupling an editable resource hub (static site) with minimal backend operations for communication/dissemination of updates and a transparent contribution system.

The portal’s architecture is intentionally shaped by the realities of the academic world, where long-term continuity is rarely guaranteed and infrastructure must survive graduate-student timelines, shifting priorities, and uneven technical support. Rather than assuming dedicated maintainers or physical servers, the design treats turnover and limited operations time as baseline through automating most routine tasks and minimizing the number of components that can silently drift or fail. Most contributions are made using standard low-barrier tools (e.g., Markdown, GitHub), while automatic deployment removes the need for direct server access or manual releases. Crucially, the system draws a hard boundary between knowledge (static, reproducible content) and operations (a small set of backend services), keeping the latter narrowly scoped and implemented as event-triggered serverless architectures that avoid always-on servers. This design turns maintenance into primarily editorial oversight, making the technical gap between a regular contributor and a maintainer practically inexistent, so anyone can take on any of these roles as required.

Beyond consolidating existing material, this platform also changes how Martini knowledge is produced and validated over time through collaborative development. In practice, all GitHub discussion threads become accessible “micro-publications” that capture the scientific rationale behind parameter choices, tutorial updates, and recommended workflows, creating an auditable trail that is difficult to reconstruct from scattered repositories and private exchanges. This centralization model also introduces added flexibilities for consistent content metadata and stronger curation layers (e.g., tagging resources by maturity, validation status, or Martini version compatibility). A natural next step would be now to formalize these criteria, so that “what is current”, “what is reliable” and what counts as “recommended” become explicit, transparent properties rather than tacit community knowledge. Such infrastructure also has the potential to shorten the iteration cycle between developers and users. For example, issues raised through help channels can be linked to solutions faster, while keeping clear authorship and version history. In this sense, the portal is less a website than a coordination mechanism, as it allows collaboration in a form that can scale naturally with community growth.

Several trade-offs remain, however, and motivate clear future directions. Importantly, pull-request review exists as the primary quality-control mechanism, but its success depends on both topic expertise and reviewer availability; when capacity is limited, review latency itself becomes a barrier to contribution. A practical response is to formalize editorial roles (e.g., topic editors for lipids, proteins, tools, etc), set explicit review expectations, and adopt consensus contribution guidelines that prioritize content organization, link integrity, meaningful cross-linking between pages and formatting styles. These changes largely require templates and agreed conventions rather than new infrastructure and will make contributor/reviewer decision-making more straightforward. In parallel, while serverless infrastructure reduces day-to-day administration, it introduces reliance on a cloud provider and requires durable storage of credentials, domain ownership, and a small amount of operational knowledge. Having these points clearly identified is a crucial first step toward finding more innovative solutions in the future.

Future work will focus on strengthening reproducibility and interoperability while keeping the barrier to contribution low. An essential step will be the integration of the Martini Database (MAD) into the portal as the canonical distribution layer for FF parameters [28, 29], enabling citable, versioned releases. Such interoperability would allow parameters to remain stored in MAD as a single source of truth with machine-readable metadata (e.g., scope, biomolecular class, version tracking, benchmarking/validation context, Martini version compatibility), while automatically synchronized with the corresponding listing entries in the portal’s *Downloads* section.

In summary, the MFFI web portal presents a pragmatic model for sustaining collaboration in academic scientific ecosystems, making community knowledge easy to contribute (docs-as-code), easy to trust (expert review), easy to maintain (automatic front-end deployment + serverless backend), and easy to disseminate (event-triggered announcements). The broader implication is that community infrastructure can be treated as a scientific instrument in its own right, one that measurably influences adoption, reproducibility, and the rate at which scientific advances propagate through a field.

## 4 Methods

### 4.1 Design requirements and guiding principles

To meet the five practical requirements initially identified (avoid reliance on a single server; minimize long-term maintenance overhead; reduce technical barriers to contribution; provide a unified home for resources; and disseminate updates in a timely manner), the portal is built around three guiding principles. First, the *docs-as-code* model enables web content to be written in plain text and tracked, reviewed, and discussed through GitHub. Second, *static-by-default* keeps the public portal as a static site, minimizing operational complexity and reducing the security surface area. Third, an *infrastructure-as-code* approach provides dynamic functionality through event-triggered AWS services. Together, these principles define a three-layer implementation: (i) a content layer authored in Markdown/Quarto; (ii) a build-and-deployment layer that renders and publishes the static site through continuous integration; and (iii) a minimal backend layer for contact submissions and opt-in email notifications. This separation keeps routine operation primarily editorial, while confining infrastructure concerns to a small set of automated services.

### 4.2 Static site generation with Quarto

The public site is generated with Quarto from `.qmd` sources into a static HTML/CSS bundle [24]. Quarto was selected because it supports: (i) low-barrier authoring via Markdown; (ii) reproducible, literate-programming style integration of code blocks and executable examples; and (iii) deterministic rendering suitable for GitHub Actions deployment.

### 4.3 Build and deployment with continuous integration workflows

The build-and-deployment layer is implemented using GitHub Actions and can be triggered either manually (`workflow_dispatch`) or automatically on every push to the `main` branch [26]. Two complementary workflows ensure that the public static site is rebuilt and published reproducibly, and that announcement posts are synchronized to the backend content library used for email distribution.

- (Static site rendering and publication) This workflow checks out the repository, installs Quarto in a clean Ubuntu runner, renders the portal, and publishes the generated static bundle to the `gh-pages` branch using the official Quarto GitHub Action. Publishing is authenticated with the repository-scoped `GITHUB_TOKEN`, enabling automated updates without manual server access and ensuring the live portal can be regenerated deterministically from the version-controlled sources on each merge to `main`.
- (Announcement source synchronization) A second workflow synchronizes announcement posts to an S3 bucket (`cgmartini-library`) whenever `main` is updated. The workflow configures AWS credentials via GitHub Secrets and uses `aws s3 sync` to upload only the Quarto source files for announcements (pattern `docs/announcements/posts/*/*.qmd`). Restricting synchronization to source files preserves a single, canonical input for both website rendering and email delivery, while S3 object-creation events under this prefix serve as the trigger for the announcement distribution pipeline described below.

### 4.4 Backend services as AWS serverless cloudformations

Backend functionality is implemented on AWS using the AWS Cloud Development Kit (CDK) [30], which synthesizes to CloudFormation templates. Backend elements include a shared AWS S3 content library (`cgmartini-library`) used to host large portal assets and two serverless stacks: (i) `AnnouncementsStack` for subscription management and announcement email delivery, and (ii) `ContactsStack` for contact form submission and message forwarding.

#### 4.4.1 Centralized file distribution via an S3 content library

To keep the front end lightweight, all heavy files are stored in a dedicated AWS S3 bucket that acts as a centralized content library. This includes downloadable parameter bundles, tutorial datasets, tools, and the announcement posts synchronized from the repository. Hosting these assets in S3 provides durable, scalable storage with stable object URLs. In addition, using S3 as the backend library enables naturally integrated automations (e.g., object-creation triggers for announcement delivery) and simplifies long-term monitoring operations.

#### 4.4.2 AnnouncementsStack: subscription management and announcement delivery

The announcements backend provides a self-managed mailing list with explicit verification and a mechanism to send email notifications as new announcements are published. Subscriptions are stored in a DynamoDB table `Subscribers` keyed by `email`. A second DynamoDB table, `VerificationTable` (also keyed by `email`), supports the verification workflow (pending/confirmed status and verification token bookkeeping). Both tables are configured with retention policies to prevent accidental deletion of subscriber state during stack updates.

A REST API, implemented with AWS API Gateway, exposes three endpoints: (i) `POST /subscribe` to initiate subscription, (ii) `GET /verifyEmail` to confirm a pending subscription, and (iii) `GET /unsubscribe` to remove an address from the mailing list. Upon subscription, the system sends a verification email via AWS Simple Email Service (SES) from, including a confirmation link that resolves to `/verifyEmail` and an unsubscribe link pointing to `/unsubscribe`.

After announcement posts are synchronized to the `cgmartini-library` bucket, an `s3:ObjectCreated:*` notification triggers a Lambda function that retrieves the post content, queries the `Subscribers` table for verified subscribers, and sends individualized emails via SES. Because distribution is triggered only by object-creation events, the system requires no continuously running services and incurs minimal steady-state overhead.

#### 4.4.3 ContactsStack: contact intake, buffering, and delivery

The contact backend implements an event-triggered contact pipeline that separates fast user-facing submission from asynchronous delivery. An API Gateway REST API exposes a `POST /contacts/new` endpoint integrated with a Lambda function (`CreateContactLambda`). On submission, the function validates each request, assigns a unique `contactId`, and stores the message in a DynamoDB table `Contacts` for durability and auditability. It then enqueues a lightweight job onto an Simple Queue Service (SQS) (`contact-processing-queue`), which decouples user-facing latency from downstream processing and absorbs bursty traffic without manual scaling.

A second Lambda (`ProcessContactLambda`) is subscribed to the queue via an event source mapping, consumes messages in batches, and sends notification emails through SES to a configured administrative recipient within the Martini developers team. Queue visibility timeouts and Lambda retry semantics provide resilience to transient failures while preserving an operational trace in CloudWatch logs.

### 4.5 Security, privacy, and operational considerations

The portal is designed to minimize the collection and handling of personal data. Backend personal information is limited to contact messages submitted voluntarily by users and email addresses required for announcements, with verification and unsubscribe endpoints providing user control over subscriptions. Operationally, the public-facing site remains static, while backend functionalities are isolated to a small set of serverless components. Each Lambda function runs under its own Identity and Access Management (IAM) execution role, with permissions scoped only to the required DynamoDB tables, read access to the S3 announcement library (for delivery), and SES sending actions. This keeps the security surface area narrowly constrained and auditable through the infrastructure definitions.

### 4.6 Pay-per-request cost model

Another advantage of the serverless components is that they follow a pay-per-request cost model. In this model, charges accrue primarily when users interact with the system (e.g., submitting a contact form, subscribing/unsubscribing, verifying an email, or publishing an announcement), while idle periods incur only in minimal cost. As a result, the backend scales automatically with short-term spikes (e.g., workshop-related traffic) without manual intervention, while remaining economical and low-maintenance during all operations.

## Code and Data Availability

All source code, including portal content, GitHub automation workflows, and backend infrastructure definitions, is publicly available in the MFFI GitHub repository (https://github.com/Martini-Force-Field-Initiative/Martini-Force-Field-Initiative.github.io) and released under the MIT License.

## Acknowledgements

D.P.T. acknowledges support from the Natural Sciences and Engineering Research Council of Canada, the Canada Research Chairs Program, and the Digital Research Alliance of Canada. R.A. acknowledges support from the Interne Fondsen KU Leuven/Internal Funds KU Leuven (STG/24/070). S.J.M. acknowledges funding from the European Research Council (ERC) with the Advanced grant 101053661 COMP-O-CELL. P.C.T.S. and L.B.A. would like to thank the support of the French National Center for Scientific Research (CNRS). P.C.T.S. acknowledges financial support through research collaboration agreements with Sanofi. P.C.T.S. and L.B.A. also acknowledge the support of the PSMN (Pôle Scientifique de Modélisation Numérique) and the Centre Blaise Pascal’s IT test platform at ENS de Lyon (Lyon, France) for the computer facilities. The platform operates the SIDUS solution developed by Emmanuel Quemener [31].

