## Supplementary Information for "Community Web Portal for Open Collaboration in the Martini Force Field Initiative"

The Martini Force Field Initiative (MFFI) web portal repository is maintained through community pull requests. This guide provides the review-based workflow used to propose, test, and integrate updates (content and site improvements), consistent with the repository's contribution procedures.

### Local environment setup

Local preview and validation of contributions are performed with Quarto. Installation instructions are platform-specific (Windows/macOS/Linux) and can be accessed at `https://quarto.org/docs/get-started/`.

Contributions can then be submitted from a personal fork as follows:

1. Fork the repository on GitHub.
2. Clone the fork locally and enter the repository directory.
3. Render the site locally:
   ```
   quarto preview --port 4040
   ```
4. Check and validate changes by visiting `http://localhost:4040`.
5. Push changes to your personal fork.
6. Open a pull request in the portal GitHub repository.

### Contribution types

The repository defines six primary contribution categories, each associated with a standard template-based workflows:

- **Publications:**
  1. Copy `docs/publications/entry_template.qmd` into the year-specific directory under `docs/publications/entries/`.
  2. Rename using a unique identifier (e.g., `author-first_word_in_title.qmd`).
  3. Complete the template fields using best-practice Markdown.
- **Announcements:**
  1. Copy `docs/announcements/entry_template.qmd` into `docs/announcements/posts/`.
  2. Create a new post folder named `YYYY-MM-DD-keywords/`. The recommended post filename is `index.qmd`.
  3. Fill in the template with the announcement details.
- **Tutorials:**
  1. Use an existing tutorial as a structural reference (e.g., any `index.qmd` under `docs/tutorials/Martini3/`).
  2. Create a dedicated tutorial directory under `docs/tutorials/Martini3/` and place all associated files (images, data) inside it.
  3. Register the tutorial by adding an entry to `docs/tutorials/Martini3/tutorials.qmd`, matching the syntax used by existing entries.
- **Force field parameters**:
  1. Add a description of the parameter set in the appropriate `.qmd` file under `docs/downloads/force-field-parameters/martini3/`, selecting the molecular class category that best matches the contribution.

2. Keep the corresponding `.itp` files available to share during review; upon approval, files can be incorporated into the backend Martini library to become downloadable through the portal.

- **Tools:**
  Tool listings are maintained within the Quarto sources under `docs/downloads/tools/`. Contributors add a description of their tool to the relevant existing `.qmd` file. Current subsections include Topology/Structure generation, Multiscaling, and Analysis.
- **General website information (e.g., list of contributors, FAQs):**
  Contributions that improve general information follow the same core workflow: implement changes, preview locally, and submit a pull request for review.

**Pull request submission workflow**

Once local validation of changes is complete, contributors can submit a pull request from their fork:

1. Commit changes:

```
git add .
git commit -m "Brief description of your changes"
```

2. Push the working branch:

```
git push origin your-branch-name
```

3. Open a pull request via the GitHub interface (Contribute → Open pull request) and complete the pull request template describing the changes.

Pull requests are assessed by designated expert reviewers. Review may result in requested revisions or direct approval. After approval, changes are merged into the main branch and the website is automatically updated and deployed.
